# Development of a *Vibrio natriegens*-based plate-clearing assay for rapid screening of PET-hydrolyzing enzymes

**DOI:** 10.1101/2025.04.02.646774

**Authors:** Susanne H. Troøyen, Ingrid M. Bolstad, Nora Drøivoldsmo, Davide Luciano, Evamaria I. Petersen, Gaston Courtade

## Abstract

Polyethylene terephthalate (PET) is a major contributor to plastic waste. Enzymatic PET degradation offers a sustainable recycling approach, but screening for effective enzymes remains challenging. Herein, we established an experimental plate-clearing assay using *Vibrio natriegens*, exploiting its protein secretion system to rapidly identify catalytic activity without time-consuming downstream processing. To validate the assay’s robustness, we tested variants of *Fusarium solani pisi* cutinase (FsC) and found that one variant, T45P, exhibited threefold higher PET-film degradation yield than the wild type and favored production of terephthalic acid. As a further test case, we screened an error-prone PCR library of FsC using the assay, obtaining a 25% hit rate after screening 150 colonies. This demonstrates the scalability of the method to large enzyme libraries.

## Introduction

Plastic materials have revolutionized modern life with their remarkable versatility and durability. However, the high chemical stability of plastics is a double-edged sword; while longevity is beneficial in many applications, their persistence in the environment has resulted in a global plastic crisis [1]. A major contributor is polyethylene terephthalate (PET), a semi-aromatic polymer of terephthalic acid (TPA) and ethylene glycol (EG) [2, 3]. Despite large-scale infrastructure for plastic production, long-term strategies for end-of-life disposal are lacking. Since landfilling and incineration pose environmental and health risks, developing efficient and sustainable methods for recycling plastic materials is increasingly important [4]. PET can be recycled mechanically, chemically, or enzymatically [5]. Mechanical recycling, the main route today, reduces material quality after each cycle. Enzymatic PET recycling has emerged as a sustainable alternative, degrading PET into a mixture of its hydrolysis products, bis(2-hydroxyethyl) terephthalate (BHET) and mono(2-hydroxyethyl) terephthalic acid (MHET), and the monomers TPA and EG. The monomers can then be repurposed for new PET production, facilitating a closed-loop recycling system that reduces the need for virgin PET production [6–9]. This not only alleviates the environmental burden of fossil-based resource consumption but also aligns with circular-economy principles by offering economic incentives through resource repurposing.

Several naturally occurring enzymes harboring PET-hydrolyzing ability have been discovered in recent years [10]. Cutinases (EC 3.1.1.74), which naturally degrade the plant polyester cutin, have attracted considerable scientific attention because many exhibit relatively high catalytic activity towards PET [8–11]. Additionally, cutinases are already widely used across several industries, including oil and dairy production [2]. One of the most successful PET-hydrolyzing enzymes to date is LCC-ICCG [12], a more thermostable variant of leaf-branch compost cutinase (LCC) [13]. Recently, a further improved version of this variant (LCC-ICCG-C09) was presented [14]. Cutinases with a Ser-Asp-His catalytic triad, like LCC-ICCG, hydrolyze PET through a nucleophilic attack on the ester carbonyl carbon by the catalytic serine, releasing BHET, MHET, TPA, and EG (Figure 1) [15, 16]. Surrounding residues aid in the mechanism. For example, a commonly conserved tryptophan residue (W190 in LCC-ICCG) facilitates binding by stacking with the benzene ring of PET, and backbone amide protons form hydrogen bonds to the carbonyl oxygen, which both increase the electrophilicity of the carbonyl carbon and stabilize the negative charge of the intermediate by forming an oxyanion hole [16, 17].

**Figure 1:**
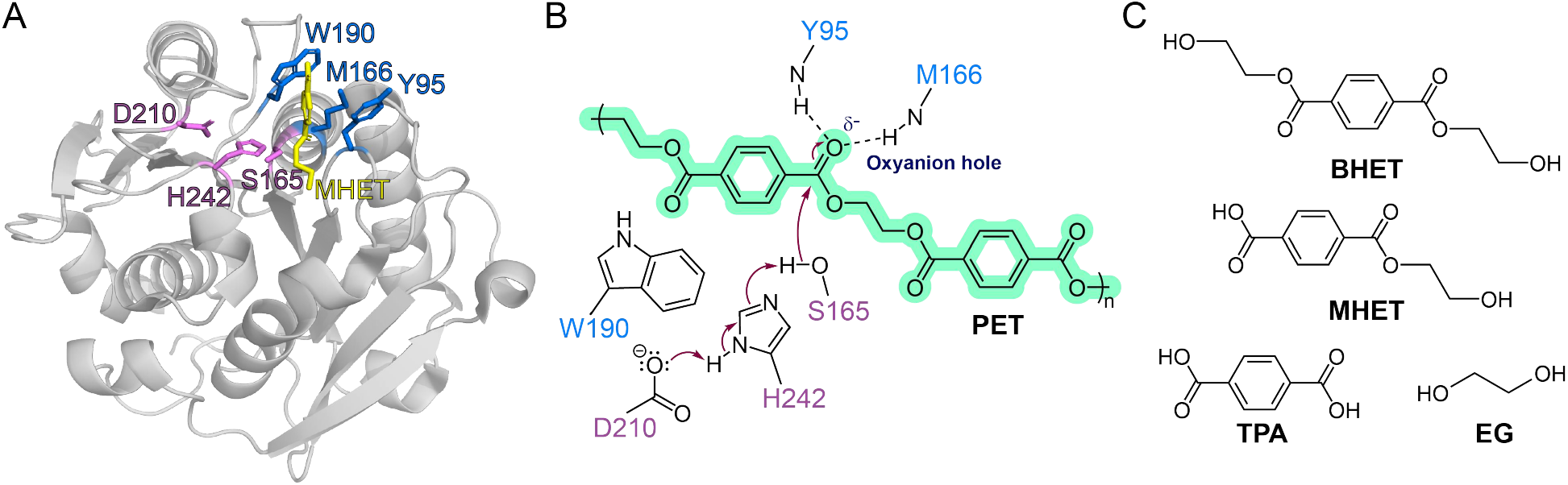
**A:** LCC-ICCG in complex with MHET (yellow sticks), based on the crystal structure of the catalytically inactive S165A variant (PDB: 7VVE [34]). Here, the catalytic triad (S165, D210, and H242) is shown in pink and the substrate-binding residues (Y95, M166, and W190) in blue. **B:** Mechanism of PET hydrolysis by LCC-ICCG, following the mechanism proposed for *Is*PETase by Han et al. [15]. Scheme adapted from Jerves et al. [16]. **C:** Chemical structures of PET degradation products.

Despite significant scientific interest, the industrial application of PET hydrolases is not trivial. Many PET hydrolases exhibit limited catalytic activity, mainly due to low thermostability [12], product inhibition [18, 19], and low activity towards high-crystallinity PET [20, 21]. It is therefore interesting to explore strategies to improve the performance of wild-type enzymes. A key challenge in this pursuit is developing efficient screening methods to rapidly identify promising enzyme variants with enhanced PET-degrading capabilities. Traditional approaches to identifying PET degradation products often involve time-consuming protein purification steps and specialized analytical techniques such as HPLC, creating a bottleneck in the screening process, particularly when evaluating large libraries of enzyme variants [22]. Several research groups have developed diverse strategies to address this challenge. For example, Bell et al. used UHPLC to screen the activity of *Is*PETase variants derived from directed evolution directly from cell lysates [23], while Ma et al. used a cell-free expression system and screened PETase activity by UV absorbance [24]. In a related approach, Ogura et al. detected enzymatic activity in *E. coli* culture medium by measuring turbidity reduction [25]; however, this method is limited by ineffective protein secretion mechanisms in *E. coli*, which are likely dependent on cell lysis.

To enable higher-throughput screening, several plate-based assays have been developed. These assays use model substrates to enable rapid visualization of activity through clearance zones around colonies expressing PET hydrolase candidates [26, 27]. Plates infused with the commercial polyester polyurethane Impranil have been used to screen large PET hydrolase libraries [28–30]. Groseclose et al. developed a high-throughput screening method for simultaneous evaluation of enzyme solubility, thermostability, and activity of large PET hydrolase libraries [31]. In this system, solubility and enzyme concentration were measured by co-expressing GFP11-tagged PET hydrolase variants with GFP1-10, producing fluorescence only when soluble enzyme is present. Activity was then evaluated by transferring partially lysed cells onto BHET-containing plates. Similarly, Wang et al. designed a plate-clearing assay based on co-expression of PETase with colicin release protein, Kil, to promote enzyme secretion in *E. coli* [32]. However, this approach suffers from drawbacks such as Kil’s negative impact on *E. coli* survivability, and false negatives when Kil activity is suppressed. While this manuscript was in revision, Lee et al. reported an alternative approach in which the YebF secretion pathway in *E. coli* is used to secrete LCC variants onto BHET plates, maintaining colony viability and avoiding the lysis drawbacks of Kil [33].

To provide a simple alternative to the methods available when this study was conceived, we developed a low-cost, rapid BHET plate-clearing assay using *Vibrio natriegens* as the expression host. *V. natriegens* offers several distinct advantages for enzyme screening applications. Most notably, it exhibits an exceptionally fast growth rate with a doubling time of approximately 10 minutes under optimal conditions, making it the fastest-growing non-pathogenic bacterium currently known [35]. This rapid growth significantly accelerates the screening process compared to traditional hosts like *E. coli*, which typically has a doubling time of 20–30 minutes. Furthermore, *V. natriegens* has previously been shown to successfully export recombinant proteins with high yields into the extracellular medium when an appropriate N-terminal export signal is fused to the protein [36, 37], a feature we considered advantageous for our plate-clearing assay. This efficient protein secretion eliminates the need for cell-lysis steps, simplifying the workflow and enabling direct assessment of enzymatic activity on substrate-supplemented plates. The *V. natriegens* strain Vmax X2 has been engineered to express genes under the control of isopropyl β-d-1-thiogalactopyranoside (IPTG)-inducible T7 promoters, allowing *E. coli* plasmids to be used directly in *V. natriegens* without the need for construction of dedicated vectors [35, 36].

We demonstrated the assay’s utility by screening four variants of *Fusarium solani pisi* cutinase (FsC), including the T45P variant described in [38], and the L182A variant described in [39, 40]. We used the benchmark cutinase LCC-ICCG as a positive control, and green fluorescent protein (eGFP) as a negative control. As the T45P variant showed high activity in the assay, we proceeded to test FsC variant activity on BHET and PET using time-resolved NMR. These experiments demonstrated the assay’s applicability in identifying promising PET hydrolase variants. We further demonstrated the scalability of the assay by applying it to an error-prone PCR mutant library of FsC. This screening platform provides a valuable tool for identifying and characterizing enzymes with enhanced PET-degrading capabilities, potentially accelerating the development of improved biocatalysts for PET recycling.

## Results and discussion

### Establishment of new plate-clearing assay

Building on previously established methods [32], we designed an efficient plate-based assay for simple detection of PET hydrolase activity. Figure 2 illustrates the assay protocol and the validation strategy. The first component to consider in such an assay is the substrate-containing agar plates. We tested three substrate-addition protocols to determine the optimal plate composition: 0.5 g/L PET powder; 0.5 g/L BHET added before and after autoclaving; and 2 g/L BHET powder added after autoclaving. Adding PET powder directly to the agar mixture was unsuitable due to its insolubility, although we note that the method presented by Charnock may be used to produce PET-supplemented agar plates [41]. BHET serves as an alternative to PET that is easier to incorporate into the agar, which we considered favorable for the practical implementation of the assay. Adding BHET before autoclaving yielded clear agar plates with no whitening, presumably because BHET was hydrolyzed during autoclaving. We found that adding 0.5 g/L BHET after autoclaving was the most suitable preparation method. It exhibited good bacterial growth and a faint, even white tint on the agar plate. Positive results were observed as clear zones around the colonies when the plates were held in front of a light source. We hypothesized that increasing BHET concentration from 0.5 g/L to 2 g/L would yield even clearer contrast in the cleared spots. However, plates with higher BHET concentrations showed impaired bacterial growth (data not shown), suggesting that this concentration was too high.

**Figure 2:**
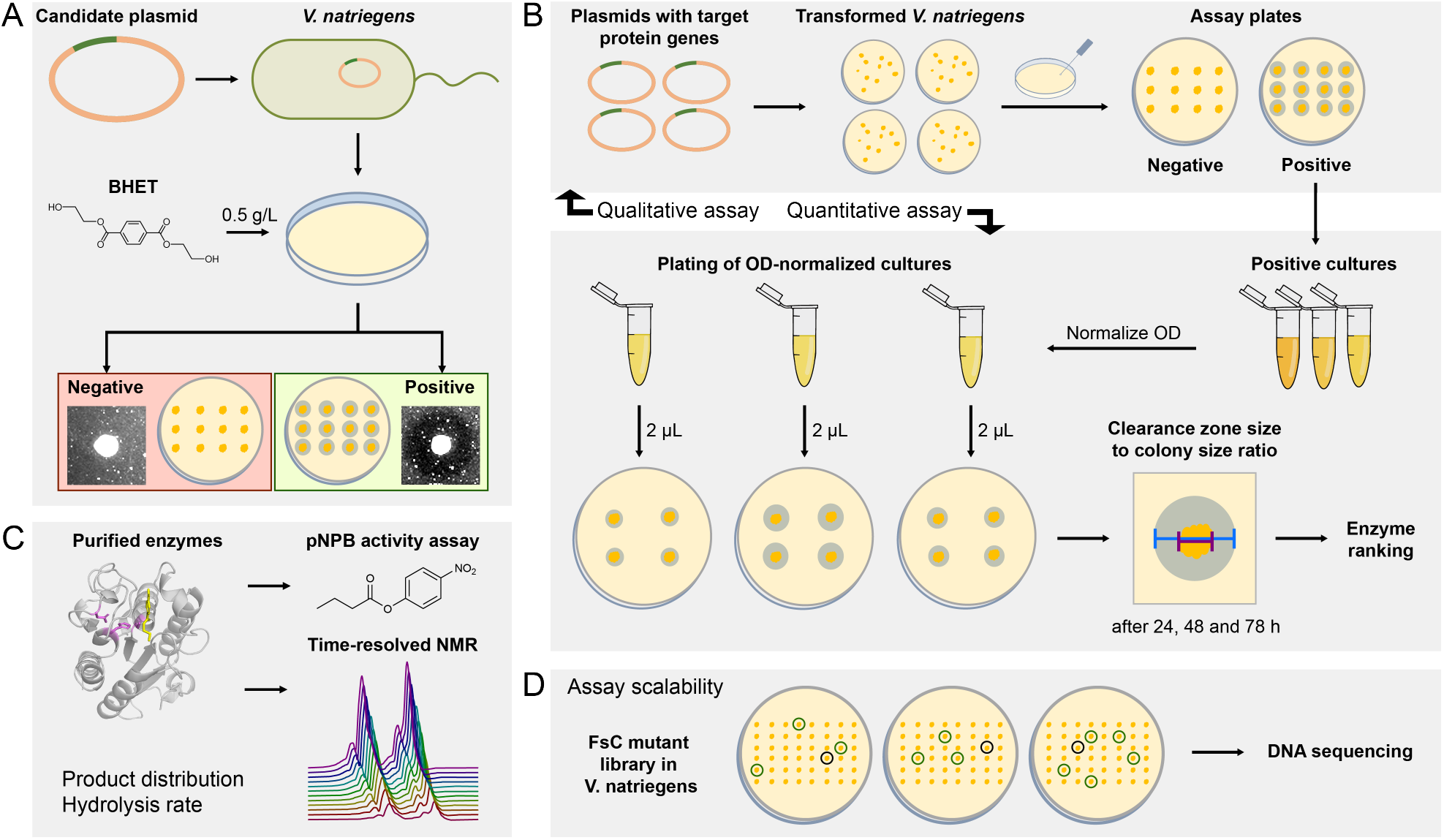
**A:** Schematic overview of the BHET plate-clearing assay employing *V. natriegens* as host for screening of PET hydrolase activity. Positive colonies display an easily observable clearance zone on the BHET plate. **B:** Protocol scheme for qualitative and quantitative plate-clearing assays, employed to test FsC variant activity, with LCC-ICCG as positive control. **C:** Illustration of experiments used to characterize hydrolytic activity of isolated FsC variants and LCC-ICCG, to demonstrate post-assay characterization of active variants. **D:** Illustration of using the assay to screen for active variants in a FsC mutant library, to verify scalability.

Next, we compared *E. coli* and *V. natriegens* as host organisms for the assay. Whereas *V. natriegens* secretes proteins into the culture medium via a type II secretion system [36], *E. coli* only translocates proteins to the periplasm. Therefore, strategies to induce periplasmic protein release are required. One such strategy involves co-expressing the target enzyme with a membrane-disrupting protein such as Kil [32]. However, this method is inherently lethal to the host organism, leading to reduced growth and productivity [42] and selecting for less active Kil variants. Despite these challenges, we attempted *E. coli* secretion for comparative purposes by introducing the *kil* gene via a bicistronic design (BCD) (Figure S1). This approach resulted in an unacceptably high rate of false negatives, rendering it ineffective as a screening tool: only 1 of 3 replicates of the positive-control LCC-ICCG showed a clearance spot (Figure S2).

In contrast, *V. natriegens* does not require specialized secretion systems, bespoke plasmids, or cell lysis before purification [35], making it an ideal host for scalable assays of PET-hydrolytic activity. Using *V. natriegens* as host showed positive results for all replicates of the positive control, LCC-ICCG, and negative results for the negative control, eGFP (Figure 3A). These results show that the BHET plate-clearing assay (Figure 2A) is effective for rapid identification of BHET-active enzymes expressed by *V. natriegens*.

**Figure 3:**
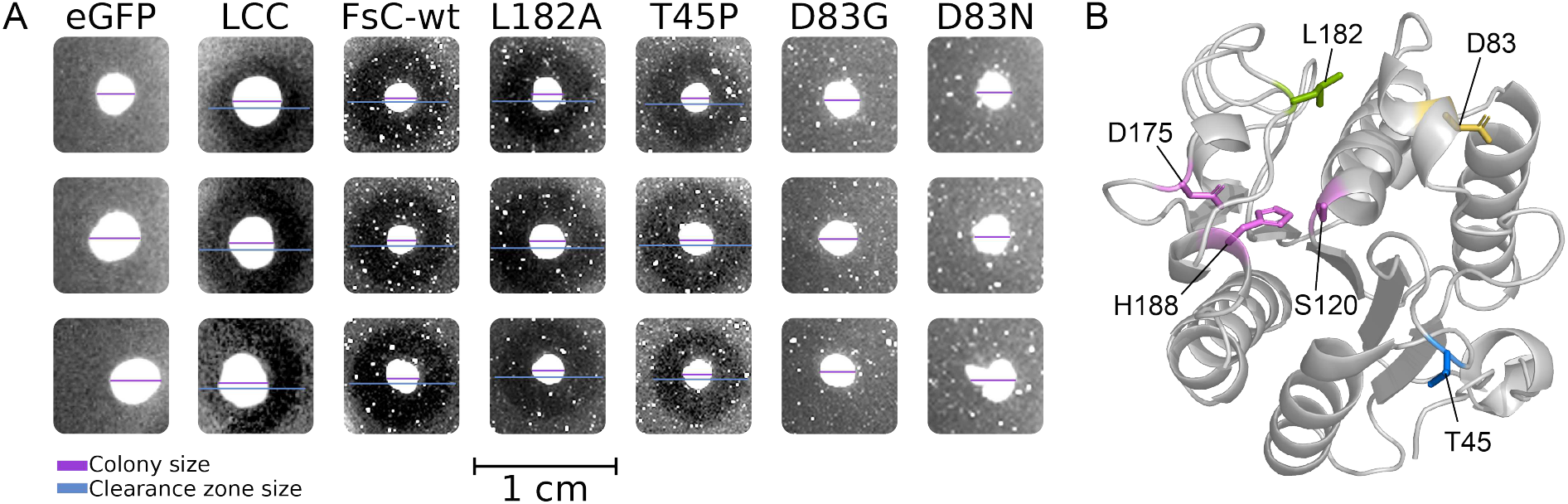
**A:** Photographs of colonies used for the qualitative plate-clearing assay in *V. natriegens* with 0.5 g/L BHET of FsC-wt, L182A, T45P, D83G, and D83N, with LCC-ICCG as a positive control and eGFP as a negative control. The photographs of the plates were cropped and color-corrected. The approximate colony diameters are indicated in purple, and the clearance zone diameters (if present) are indicated in blue. **B:** FsC crystal structure (PDB: 1CEX [47]). Mutated residues (L182, T45 and D83) and the catalytic triad (S120, D175 and H188) are shown as sticks.

### Screening of rationally selected FsC variants

To demonstrate the application of the assay to identify BHETase activity in cutinase variants, we screened FsC-wt and four variants (L182A, T45P, D83G, and D83N), using LCC-ICCG as a positive control and eGFP as a negative control. Previous studies have shown that the L182A variant is active on PET and displays higher fractions of TPA in the product mixture compared to the wild-type enzyme [39, 40]. The introduction of proline at position 45 (T45P) has been predicted to alter the distance between the E44 side chain and the catalytic S120, thereby influencing the geometry and charge distribution of the oxyanion hole [38]. Because the activity of the T45P variant on PET and BHET is unknown, we included it in our assay. The negatively charged D83 is located on a flexible helix that has been demonstrated to affect the structural substrate selectivity of FsC, as well as the integrity of the oxyanion hole [43, 44]. NMR studies have shown that D83 is involved in BHET binding [40], which prompted us to screen two variants substituting the aspartate with glycine and asparagine.

Of the screened enzymes, wild-type FsC (FsC-wt) and the L182A and T45P variants gave positive clearance zones on the BHET plates (Figure 3). In contrast, the D83G and D83N variants yielded negative results, showing that alteration of the properties of the flexible helix severely impacts hydrolytic activity on BHET.

This initial qualitative screening method was performed by transferring recombinant *V. natriegens* colonies directly to the BHET assay plates. To test a quantitative approach, we cultured the cells and normalized cell density to ensure that an equal amount of cells was applied to the plates (Figure 2B). Clearance zones around colonies were measured after 24, 48, and 72 hours, and the ratio of the clearance zone size to the colony size was used as a high-throughput measure of activity (Figure S3). Among the enzymes tested, FsC-wt and T45P exhibited the largest clearance zones, followed by LCC-ICCG and L182A. T45P displayed faster clearance zone growth than the other variants. Consistent with the qualitative screening, D83G and D83N showed little to no plate clearance, confirming their poor activity.

Our quantitative assay provides rapid estimates of relative activity among screened variants with similar expression levels. However, the ratio of the clearance zone size to the colony size cannot serve as a definitive measure of enzymatic activity because it does not account for enzyme expression levels. In addition, the assay is temperature-limited to accommodate *V. natriegens* growth (30°C), which could fail to detect BHETase activity in highly thermostable enzymes. Despite these limitations, the assay provides a rapid method for screening large pools of PET hydrolase candidates.

Some cutinases, including LCC-ICCG and *Thermobifida fusca* cutinase (TfC), have been shown to permeate cell membranes upon expression [45, 46] so plate-clearing assays do not necessarily require a dedicated secretion system for such enzymes. However, enzymes that lack this ability must be expressed in a system that facilitates secretion, for example by fusion to secretion proteins such as Kil [32] or YebF [33]. The *V. natriegens* assay offers two main advantages over *E. coli* -based assays. Firstly, it eliminates the need to add secretion tags and to remove them for post-assay analysis of positive enzymes, as *V. natriegens* naturally secretes enzymes into the medium. Secondly, the fast growth rate of *V. natriegens* enables faster visualization of clearance zones (1 day) and more efficient protein expression and purification compared to using *E. coli* [36].

### Hydrolytic activity of purified FsC variants

The quantitative plate-clearing assays indicated that T45P was the most active FsC variant, prompting further characterization of its enzymatic activity relative to FsC-wt, L182A, and LCC-ICCG (Figure 2C). We evaluated BHET degradation using time-resolved ^1^H-NMR over 1 hour (Figure 4A). Analysis of the initial rates showed that LCC-ICCG was the fastest BHETase, degrading 75% of the substrate within the first 20 minutes. T45P exhibited slightly lower BHETase activity to FsC-wt, while L182A displayed a 3-fold reduction in BHET degradation rate.

**Figure 4:**
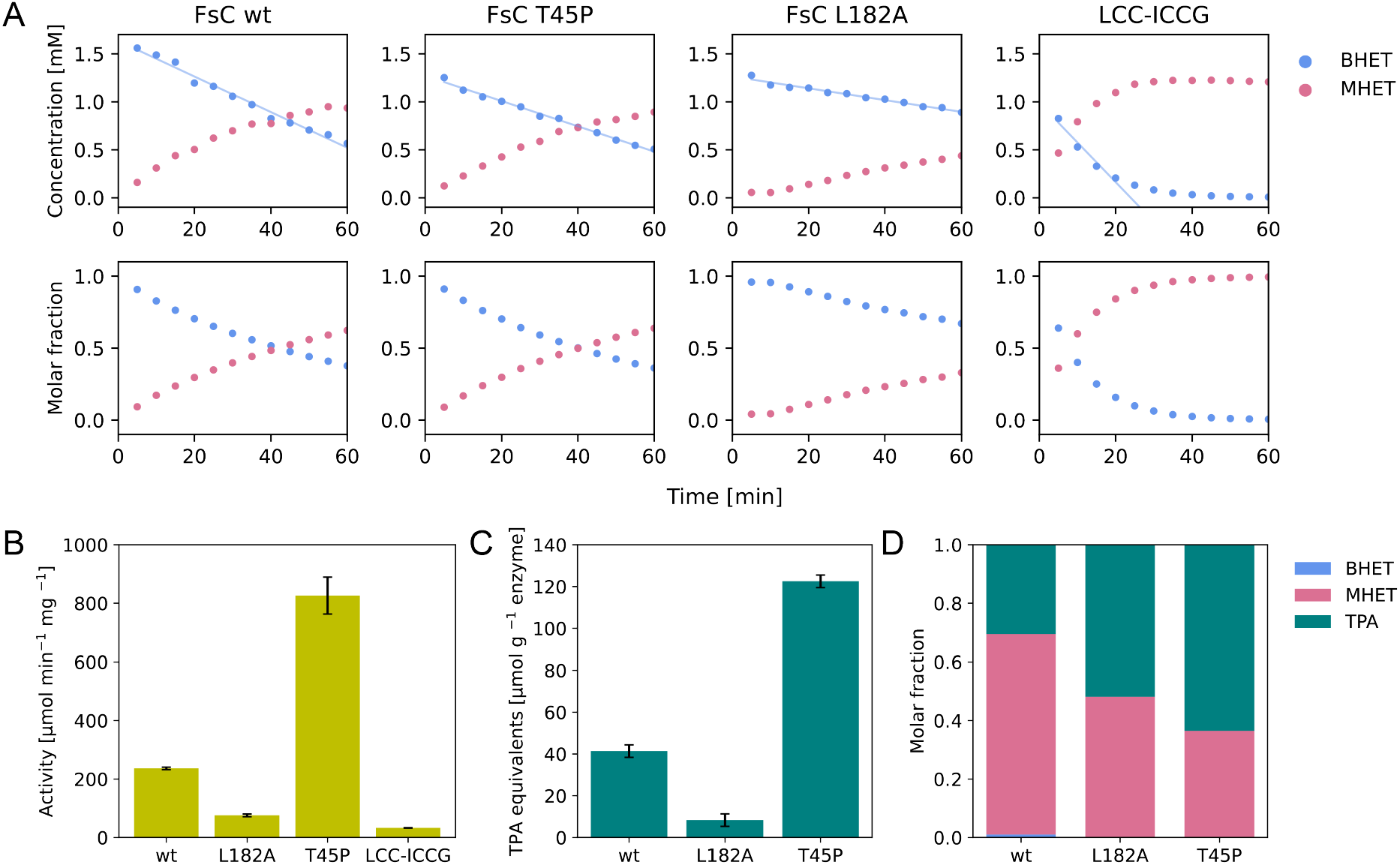
Catalytic activity of PET-degrading enzymes **A:** Time courses of BHET hydrolysis by FsC-wt, T45P, L182A, and LCC-ICCG measured by time-resolved ^1^H-NMR at 40°C. Spectra were recorded every 5 minutes for 60 minutes. The top panels show BHET (blue) and MHET (pink) concentrations over time, calculated from their ratios relative to a standard TSP signal at 400 µM. The bottom panels show BHET and MHET molar fractions over time. Initial BHET degradation rates (blue lines) were determined to 13.9 ± 1.2 (FsC-wt), 9.9 ± 0.6 (T45P), 4.6 ± 0.5 (L182A), and 24.7 ± 7.4 (LCC-ICCG) µmol g^-1^ enzyme s^-1^. **B:** Hydrolytic activity on pNPB at 20°C determined by spectrophotometry, expressed as the rate of 4-nitrophenol formation (µmol min^-1^) per mg of enzyme in a total volume of 1 mL. Error bars represent the standard deviation from triplicate experiments. **C:** Total product yield (in TPA equivalents) after 16.3 hours of PET degradation at 40°C, determined by NMR. Error bars represent the standard deviation in the TSP signal between the samples. **D:** Relative ratios of BHET (blue), MHET (pink), and TPA (green) after 16.3 hours of PET degradation at 40°C, determined by NMR. In **C** and **D**, data for FsC-wt and L182A were previously published [40].

To determine whether the differences in BHETase activity reflected broader changes in catalytic efficiency, we also evaluated general esterase activity using 4-nitrophenyl butyrate (pNPB) as a model substrate in spectrophotometric absorbance assays. The pNPB assay revealed a 3.5-fold activity increase for T45P and a 3-fold reduction for L182A, compared to FsC-wt (Figure 4B). In contrast to the NMR results with BHET, LCC-ICCG showed the lowest activity among all tested enzymes on pNPB.

To evaluate PET degradation activity, we determined product yields of the T45P variant after 16.3 hours of incubation with PET film by ^1^H-NMR. Full time-resolved product distributions from this reaction are shown in Figure S4. The results were compared with previously published data for FsC-wt and L182A [40] (Figure 4C and 4D). At 120 µmol TPA equivalents per gram of PET, T45P outperformed both FsC-wt (40 µmol/g) and L182A (10 µmol/g), demonstrating a 3-fold higher product yield than the wild-type enzyme (Figure 4C). Additionally, T45P produced approximately 2-fold higher TPA-to-MHET ratio than FsC-wt (Figure 4D), underscoring its enhanced efficiency in converting MHET to TPA.

In all, the trends in hydrolytic activity on BHET, pNPB, and PET film were consistent with the growth of the BHET clearance zones observed in the quantitative plate-clearing assay (Figure S3), except for LCC-ICCG, which demonstrated superior BHET degradation in the NMR assay. We suggest that low enzyme production levels may contribute to the lower clearing rate observed for LCC-ICCG in the quantitative plate-clearing assay. However, the temperature dependence of hydrolysis rates should also be considered, as the NMR assay was performed at 40°C, whereas the plate-clearing assay was carried out at 30°C for optimal *V. natriegens* growth. Previous pNPB-based hydrolysis assays have shown an optimal temperature of around 20–25°C for FsC-wt, with activity nearly halved at 35°C [38]. Conversely, LCC-ICCG has been shown to exhibit lower activity at low temperatures and a higher reported temperature optimum of around 50–70°C [48]. This is consistent with our pNPB assay (Figure 4B), which showed significantly lower esterase activity for LCC-ICCG than for FsC-wt at 20°C.

Our results show that T45P is a more efficient PETase than FsC-wt and L182A, particularly because it prevents MHET accumulation, a known inhibitor of PET-degrading enzymes [19]. While the structural alteration from the T45P substitution likely contributes to the reduced MHET accumulation, the exact mechanistic basis for this improvement remains unclear. Beyond its catalytic efficiency, T45P exhibits additional traits that enhance its industrial potential. Previous studies have reported increased thermostability and a broader pH-activity profile for T45P compared to FsC-wt [38], both of which are highly desirable for large-scale applications. Although a detailed mechanistic investigation is beyond the scope of this work, T45P represents a compelling case study for exploring how active-site geometry and electrostatic environment influence the distribution of hydrolysis products during PET degradation.

### Large-scale screening of FsC mutant library

To evaluate the scalability and robustness of the plate-clearing assay, we transformed a randomly mutagenized FsC library generated via error-prone PCR into *V. natriegens* (Figure 2D). A plate containing several hundred colonies was obtained, from which 150 colonies were transferred onto six BHET assay plates to screen for enzymatic activity (Figure S5). Of the 150 screened colonies, 38 mutants exhibited clearance zones, indicating catalytic activity.

For sequence analysis, all active mutants, along with six inactive mutants (one from each plate), were subjected to Sanger sequencing. Sequencing revealed that 27 of the 38 active variants carried wild-type amino acid sequences, suggesting either silent mutations or no mutations at all. The remaining 11 active variants carried substitutions (Table S1): nine harbored single-amino-acid substitutions, and two contained two substitutions each. Among the four successfully sequenced inactive variants, one carried a single substitution, one a double, one a triple, and one a quadruple substitution.

Notably, 7 of the 13 substitutions identified in active mutants were non-conservative according to the BLOSUM62 substitution matrix [49], highlighting FsC’s tolerance to structurally disruptive amino acid substitutions while retaining catalytic activity. In contrast, inactive variants exhibited a higher substitution load; only one substitution was conservative, and most variants carried multiple substitutions per enzyme. To assess the structural implications of these substitutions, we mapped them onto the three-dimensional structure of FsC (Figure 5). The analysis revealed a striking spatial distribution. Substitutions in active variants were primarily located away from the active site, suggesting that peripheral structural alterations can be tolerated without abolishing function. In contrast, substitutions in inactive variants were closer to the active site, with two substitutions (D175G and H188P) occurring directly within the catalytic triad.

**Figure 5:**
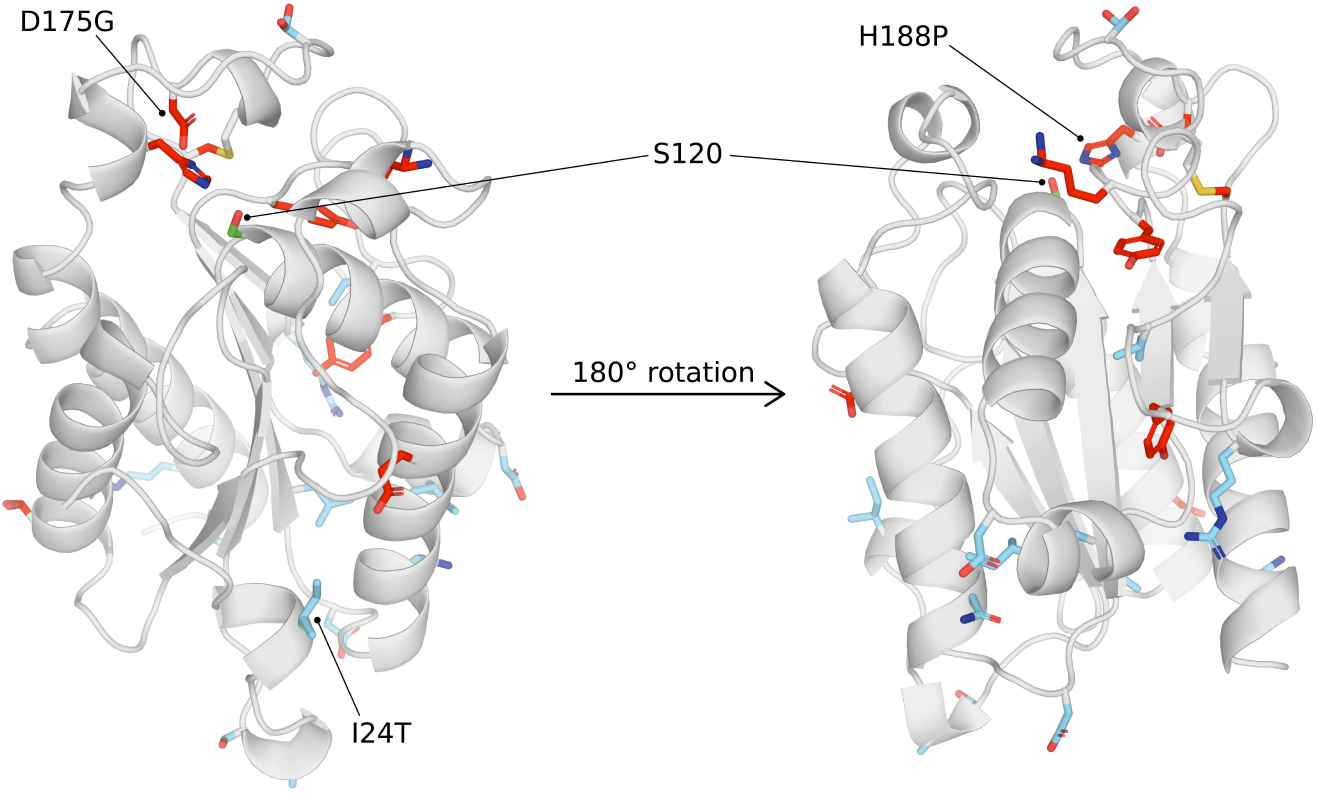
3D representations of FsC (PDB: 1CEX [47]) displayed in two different perspectives. Amino acid mutations identified in active variants are shown in blue, whereas those for the inactive variants are shown in red. The three amino acids that form the catalytic triad (S120, D175, H188) in the active site of FsC are labeled. S120 is shown as green sticks, as this residue was not substituted in any of the screened variants. The active variant displaying the largest clearance zone (I24T) is also labeled. The figure was created using PyMOL [50].

To evaluate the relationship between clearance zone size and enzymatic activity in our library, we produced and purified the FsC variant showing a particularly large clearance zone-to-colony size ratio, FsC-I24T (colony D16 in Figure S5). The non-conservative I24T substitution is located far from the active site, on the protein surface (see Figures 5 and S6). The BHET hydrolysis rate of this variant was identical to that of the wild-type enzyme (see Figure S7), indicating that the substitution does not affect catalytic activity. This finding demonstrates that larger clearance zones do not necessarily translate into higher enzymatic activity and instead suggests that this variant is expressed at a higher level than the wild type. However, substitution of the hydrophobic isoleucine with a polar threonine on the protein surface could increase structural stability around the C-terminus, possibly through hydrogen bonding with nearby glutamine residues (Figure S6). These substitutions away from the active site may influence the thermostability, as observed for LCC-ICCG in previous studies [14], and could be of interest for future studies on engineering FsC.

By screening 150 colonies transformed with FsC mutant library DNA, we show how the plate-clearing assay can screen hundreds of colonies in one day without specialized equipment. However, demand for higher-throughput screening methods is high as the pool of enzymes with potential PET hydrolase activity rapidly expands [27]. The throughput of this method is limited mainly by the time required to transfer individual colonies from master plates to assay plates. To increase throughput, more efficient colony transfer methods should be used, such as spreading mixed transformant suspensions [28, 30] or transferring colonies via semi-permeable membranes [31]. Additionally, we incubated the assay plates at 30°C, which is significantly lower than the desired working temperature of industrially relevant PET hydrolases. Thermostable enzymes with higher optimal temperatures may show lower clearance rates at 30°C and be falsely discarded as low-activity candidates. Branson *et al*. have shown that plasmids can be recovered from screening plates even after heat treatment, provided colonies are allowed to form on the plate before incubation at elevated temperatures [30]. Building on their approach, our assay may be modified to account for thermostability during screening. This simple assay may be used for discovery of new PET-hydrolyzing enzymes from metagenomic libraries or *de novo* enzyme design.

## Conclusions

We developed a simple, low-cost plate-clearing assay employing *V. natriegens* as a host organism for rapid screening of PET-hydrolyzing enzymes. We used the assay for preliminary screening of BHETase activity in rationally selected FsC variants using qualitative and quantitative approaches, demonstrating its success in uncovering two key findings. First, we identified D83G and D83N as inactive variants, highlighting the importance of the flexible helix in regulating FsC activity. Second, we purified the active variants and found greatly increased PETase activity and reduced MHET accumulation for T45P. Using a quantitative assay approach, we ranked the FsC variants according to clearance zone growth rates. NMR and pNPB assays on purified enzymes showed a similar ranking, where the trend was consistently lower activity for L182A and higher activity for T45P, compared to the wild-type FsC. However, this correspondence did not apply to LCC-ICCG, where we observed a much higher BHET degradation rate by NMR. This suggests that the quantitative assay may not be ideal for enzymes with higher optimal temperatures for PET activity, or for enzymes with low expression levels in *V. natriegens*. We found that the main advantages of the method are its simplicity, its low cost, and its rapid execution. We used the plate-clearing assay to screen an FsC mutant library, showing how hundreds of clones can be screened in a day. Here, 11 active FsC variants were identified, all carrying amino acid substitutions far from the active site.

## Materials and methods

### Plasmid construction

Plasmids for LCC-ICCG and eGFP were constructed using pET21(+) plasmids (Twist Bioscience). Target proteins were codon-optimized for *E. coli* and the signal peptide PhoA fused to the N-terminus for periplasmic export. To achieve protein secretion into the medium by *E. coli* in the *E. coli* -based plate-clearing assays, we incorporated the Kil protein (colicin release lysis protein, gene ID 2693958) into our LCC-ICCG and eGFP designs. We implemented a bicistronic design (BCD) based on Sun et al. [51] to enable simultaneous uptake and expression of both our target gene and the *kil* gene. The BCD consisted of two key elements: (1) a fore-cistron sequence containing the target protein gene with a downstream overlapping Shine-Dalgarno (SD) sequence, and (2) a TAATG sequence that served as both a stop codon (TAA) for the target gene and a start codon (ATG) for the *kil* gene (see Figure S1). For expression of FsC and FsC variants (D83G, D83N, T45P, L182A), plasmids were constructed as described previously [38]. The *kil* gene was not included in the plasmids for FsC enzymes. The same FsC mutant plasmids were used to transform *V. natriegens* and *E. coli* for plate-clearing assays and protein production, respectively.

### Transformation

*E. coli* BL21 (DE3) T7 Express (NEB catalog number 2566) strains were transformed using a standard heat-shock protocol. In short, DNA (1 ng/µL) was added to chemically competent cells and incubated on ice for 20 min, then 30 s at 42°C, then on ice for 2 min. The cells were incubated in 1 mL super optimal broth (20 g/L tryptone, 5 g/L yeast extract, 0.5 g/L NaCl, 0.186 g/L KCl, 0.952 g/L MgCl_2_, 3.6 g/L glucose) for 1 h at 37°C, 220 rpm, before plating on LB agar supplemented with 100 µg/mL ampicillin.

*V. natriegens* Vmax X2 (TelesisBio catalog number CL1300-10) strains were transformed using electroporation. Electrocompetent Vmax X2 cells were prepared using a modified procedure from Weinstock et al. [35]. 10 mL LBv2 medium (10 g/L tryptone, 5 g/L yeast extract, 5 g/L NaCl supplemented with v2 salts: 204 mM NaCl, 4.2 mM KCl, 23.14 mM MgCl_2_) was inoculated with *V. natriegens* and incubating overnight at 20°C, 200 rpm. 250 mL LBv2 was inoculated with 1% pre-culture and incubated at 30°C, 200 rpm until OD_600_ = 0.6. Cells were incubated on ice for 15 min and harvested by centrifugation at 6500*g* for 20 min at 4°C. The pellet was resuspended in 10 mL electroporation buffer (7 mM potassium phosphate, 680 mM sucrose, pH 7.0). The suspension was diluted to 35 mL with electroporation buffer, and centrifuged at 6200*g* for 15 min at 4°C. The supernatant was carefully discarded, and the cells were washed twice with 35 mL electroporation buffer followed by centrifugation at 6200*g* for 15 min at 4°C. The cells were resuspended in electroporation buffer to a final OD_600_ = 16. Cells were snap-frozen in liquid nitrogen in 50 µL aliquots and stored at -80°C. For transformation, electrocompetent Vmax X2 cells were thawed on ice for 5 min. Plasmid DNA (1 ng/µL) was added to the cells. The cells were incubated on ice for 3–5 minutes. The cell-DNA mixture was transferred to a pre-chilled electroporation cuvette with a 0.1 cm gap size. Electroporation was performed using an ELEPO21 Electroporator (Nepagene) with the following parameters: voltage: 1000 V, pulse width: 3.5 ms, pulse interval: 50 ms, no. of pulses: 1, polarity: +, followed by a poring pulse: 100 V, pulse width: 50 ms, pulse interval: 50 ms, no. of pulses: 3, polarity: +/-. The cells were then transferred to a 15 mL tube with 450 µL recovery medium (LBv2 + 680 mM sucrose). To transfer any remaining cells, the cuvette was washed with 50 µL recovery medium. The cells were recovered by incubation at 30°C, 220 rpm for 2 h, before plating 200 µL cells on LB-agar plates with 50 µg/mL ampicillin.

### Plate-clearing assays

We tested *E. coli* and *V. natriegens* as host organisms in the plate-clearing assays. Strains harboring LCC-ICCG and eGFP plasmids were used as positive and negative controls, respectively. We employed the plate-clearing assay in *V. natriegens* to screen for catalytic activity of FsC-wt, D83N, D83G, L182A, and T45P.

The plate-clearing assay method was modified from Wang et al. [32]. Recombinant colonies of *E. coli* or *V. natriegens* were transferred onto LB agar plates supplemented with ampicillin (100 µg/ml for *E. coli*, 50 µg/ml for *V. natriegens*), 0.1 mM IPTG, and substrate. For the *V. natriegens* assays, we tested three substrate compositions: 0.5 g/L PET powder (particle size < 300 µm, crystallinity >40%, Goodfellow, Huntingdon, UK, cat.no. ES30-PD-000131), 0.5 g/L BHET, and 2 g/L BHET powder (Sigma-Aldrich, cat. no. 465151, ground using a mortar). For subsequent assays, 0.5 g/L BHET plates were used. Each clone was tested in triplicate. After incubation at their respective optimal temperatures (37°C for *E. coli* and 30°C for *V. natriegens*) for 24–48 hours, plates were visually inspected for clearance zones around colonies, indicating positive enzyme activity.

For the quantitative assay approach, pre-cultures were prepared by inoculating 1 mL LBv2 medium with cells from a recombinant colony and incubating at 30°C and 220 rpm overnight. The following day, 500 µL pre-culture was transferred to 10 mL LBv2 medium and incubated at 30°C and 220 rpm. After 2.5 hours, all samples were normalized to OD_600_ = 0.8. 2 µL of each clone was added to an assay plate, with four replicate wells per clone. The plates were imaged at 24, 48, and 72 h, and the diameters of the clearance zones and colonies were measured using ImageJ [52]. At each time point, the ratio of the clearance zone to the colony diameter was calculated for each clone as the average of the four replicates.

### Screening of FsC mutant library in *V. natriegens*

The FsC mutant library was created with error-prone PCR (epPCR). The PCR was performed as described in Xu et al. [53], with manganese added in the first round and dITP in the second. Chemically competent

*E. coli* DH5α cells (NEB catalog number C2987) were transformed with the FsC mutant library DNA by heat shock, as described above. Instead of plating the cells, the whole tube of transformed cells was used to inoculate 5 mL LB supplemented with 100 µg/ml ampicillin, and the culture was incubated overnight at 37°C, 200 rpm. The DNA was isolated using a Monarch Plasmid Miniprep kit (NEB catalog number T1110) following the standard protocol. DNA was stored at -20°C. *V. natriegens* cells were transformed with library DNA by electroporation, as described above, producing ∼10^6^ colony-forming units (CFU) per µg DNA. Random colonies from the plate were picked and systematically transferred to a new agar plate containing 50 µg/mL ampicillin, 0.5 g/L BHET, and 0.1 mM IPTG. Plates were incubated at 30°C overnight and then inspected the following day. To extract DNA for sequencing, a pre-culture was prepared by inoculating 5 mL of LBv2 containing 50 µg/mL ampicillin with a pipette tip dipped into the colony. Pre-cultures were grown overnight at 30°C, and plasmid DNA was extracted from each pre-culture.

### Protein production and purification

The FsC enzymes of interest were produced in *E. coli* BL21 (DE3) following a procedure described previously [40]. LCC-ICCG was produced in *V. natriegens*. Pre-cultures were made by inoculating 5 mL enhanced 2xYT medium (20 g/L yeast extract, 32 g/L tryptone, 17 g/L NaCl, 17.6 mM sodium phosphate, 0.2% glucose, pH 7.4) supplemented with 50 µg/mL ampicillin with the recombinant *V. natriegens* strain and incubating at 30°C and 225 rpm overnight. 50 µL of overnight culture was added to 50 mL of enhanced 2xYT medium containing 50 µg/mL ampicillin in a 250 mL baffled shake flask and grown for 2 hours at 30°C and 225 rpm, until OD_600_ > 0.5. Protein expression was induced by adding IPTG to a final concentration of 1 mM. Cells were then incubated for 24 hours at 30°C and 225 rpm to produce the recombinant protein. As the LCC-ICCG plasmid was constructed with an export signal (PhoA), recombinant proteins were harvested from the growth media by centrifugation at 6000*g* for 15 minutes at 4°C. The supernatant was sterile-filtered (0.2 µm pore size) before further protein purification.

The proteins were purified with FPLC using an ÄKTA pure chromatography system (Cytiva). The periplasmic extracts were applied to a 5 mL weak anion-exchange HiTrap DEAE Sepharose FF column (Cytiva) that had been equilibrated with 20 mM Tris-HCl (pH 7.0). LCC-ICCG was purified using cation-exchange chromatography with a 5 mL HiTrap CM Sepharose FF column (Cytiva) and 20 mM HEPES (pH 7.5) as equilibration buffer. The purification protocol for all proteins was as follows: after sample application, a 5 CV wash with equilibration buffer, followed by elution with a linear NaCl gradient from 0 to 0.5 M over 50 CV, and a final wash of 3 CV with 1 M NaCl. Collected fractions displaying an A_280_ absorption peak during the elution phase were analyzed with SDS-PAGE. Fractions containing the protein of interest were concentrated using VivaSpin 20 centrifugal concentrators (Sartorius) with a molecular weight cut-off of 10 kDa at 7000*g* and 10°C for 10 minutes. The total yield of LCC-ICCG produced in *V. natriegens* and purified from the supernatant of the growth medium had a yield of ∼5 mg/L cell culture. Consistent with our previous results [40], the yields of the FsC proteins produced in *E. coli* were ∼40 mg/L.

### pNPB assay

Esterase activity assays were performed to assess the hydrolytic activity of FsC, FsC-L182A, FsC-T45P, and LCC-ICCG on 4-nitrophenyl butyrate (pNPB). The activity was determined by measuring the absorbance (UV-1800, Shimadzu) of 4-nitrophenol at 410 nm (extinction coefficient: 15548 M^-1^ cm^-1^) using pNPB as the substrate over a reaction time of 15 min. All measurements were performed at 20°C in 50 mM Tris-HCl (pH 8). The pNPB concentration was 4 mM. The activity was expressed as units per mg of enzyme. One unit is defined as the appearance of 1 µmol of 4-nitrophenol per minute in a 1 mL reaction volume. All measurements were run in triplicate and background corrected for autohydrolysis in the buffer.

### NMR spectroscopy analysis of FsC variant activity on BHET and PET

NMR spectra were recorded on a Bruker Ascend 800 MHz NMR spectrometer (Bruker BioSpin AG, Switzerland) equipped with an Avance NEO console and a 5 mm cryogenic CP-TCI z-gradient probe at the NV-NMR Center at NTNU, which is part of the Norwegian NMR Platform (NNP). Signals from BHET, MHET, TPA, and TSP were integrated in Bruker TopSpin version 4.3.0.

Time-resolved ^1^H-NMR was used to monitor BHET degradation by FsC-wt, T45P, L182A, I24T, and LCC-ICCG. NMR samples consisted of 2 mM BHET in 25 mM sodium phosphate (pH 6.5) supplemented with 10% D_2_O and 400 µM trimethylsilylpropanoic acid-d4 (TSP), prepared in 5 mm NMR tubes with a total volume of 600 µL. After addition of enzyme to a final concentration of 1 µM, the tube was immediately inserted into the spectrometer. ^1^H-NMR spectra were recorded at 40°C every 5 min over 1 h, for a total of 12 spectra.

The hydrolytic activity of FsC-T45P on PET was measured under almost identical conditions as previously characterized FsC variants [40], except for the concentration of D_2_O in the buffer, which was 10% and not 99.9%. A piece of PET film (GoodFellow product ES301445; 2.0 ± 1.6% crystallinity [18], cut to 30 × 4 × 0.25 mm) was placed in in 25 mM sodium phosphate (pH 6.5) supplemented with 50 mM NaCl, 10% D_2_O, and 400 µM trimethylsilylpropanoic acid-d4 (TSP), in a 5 mm NMR tube at a total volume of 600 µL. After addition of FsC-T45P to a final concentration of 10 µM, the sample was immediately inserted into the spectrometer. ^1^H-NMR spectra were recorded at 40°C every 5 min over 16.3 h, for a total of 196 spectra.

## Data availability

Data and Python scripts used for data processing and making the figures are available from https://github.com/gcourtade/papers/blob/master/2025/Vnat_denovo.

## Acknowledgements

G.C. gratefully acknowledges funding from the Novo Nordisk Foundation (grant number NNF22OC0073963) and support from the Norwegian NMR Platform (NNP; grant number 226244 from the Research Council of Norway). We are grateful to Prof. Peter Fojan for his valuable insights and productive scientific discussions.

## Supporting information

- SI: Supplementary Information Figure S1–S7; Table S1

