## Supplementary Information for "Development of a *Vibrio natriegens*-based plate-clearing assay for rapid screening of PET-hydrolyzing enzymes"

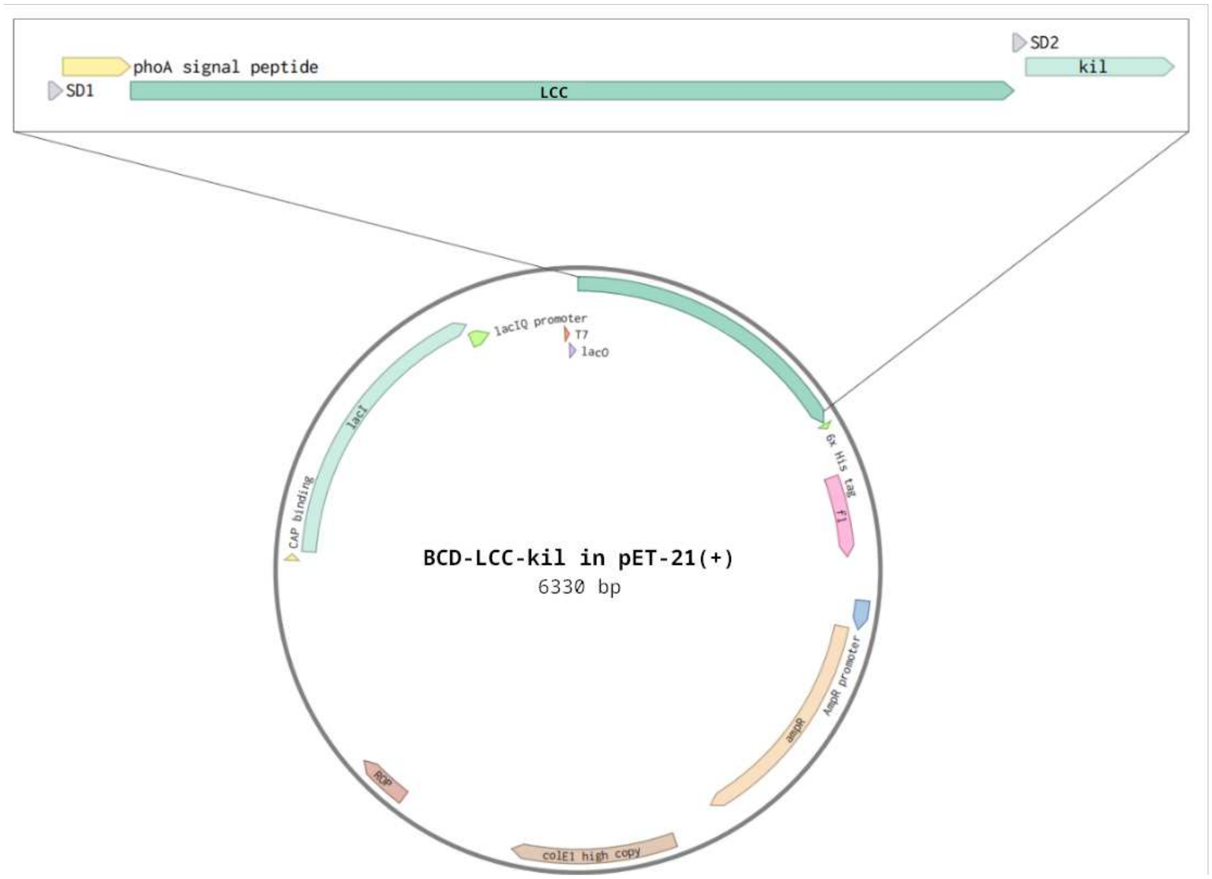


Figure S1: Schematic of BCD-LCC-kil in pET-21(+). The inset shows the bicistronic design (BCD) downstream of the T7 promoter. The first Shine-Dalgarno sequence (SD1; AAGGAGATATACC) directs translation of LCC-ICCG with a phoA signal peptide for translocation to the periplasm, whereas the second Shine-Dalgarno sequence (SD2; AGGAGATATATAA), embedded at the 3' end of the LCC coding sequence, directs translation of *kil* (colicin release lysis protein; gene ID 2693958).

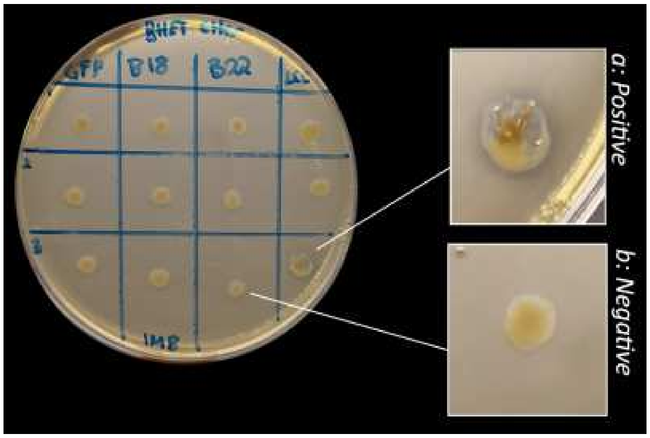


Figure S2: Photograph of a plate-clearing assay on a plate supplemented with 0.5 g/L BHET after autoclaving. Three colonies of eGFP and LCC-ICCG are shown. B18 and B22 are shown for completeness but were not considered further in the analysis. The photograph was taken after parts of the positive colony had been removed. a: Close-up view of the LCC-ICCG colony with a surrounding clearance zone (positive clearing). b: Close-up view of a colony without a clearance zone (negative clearing). All other colonies gave negative results.

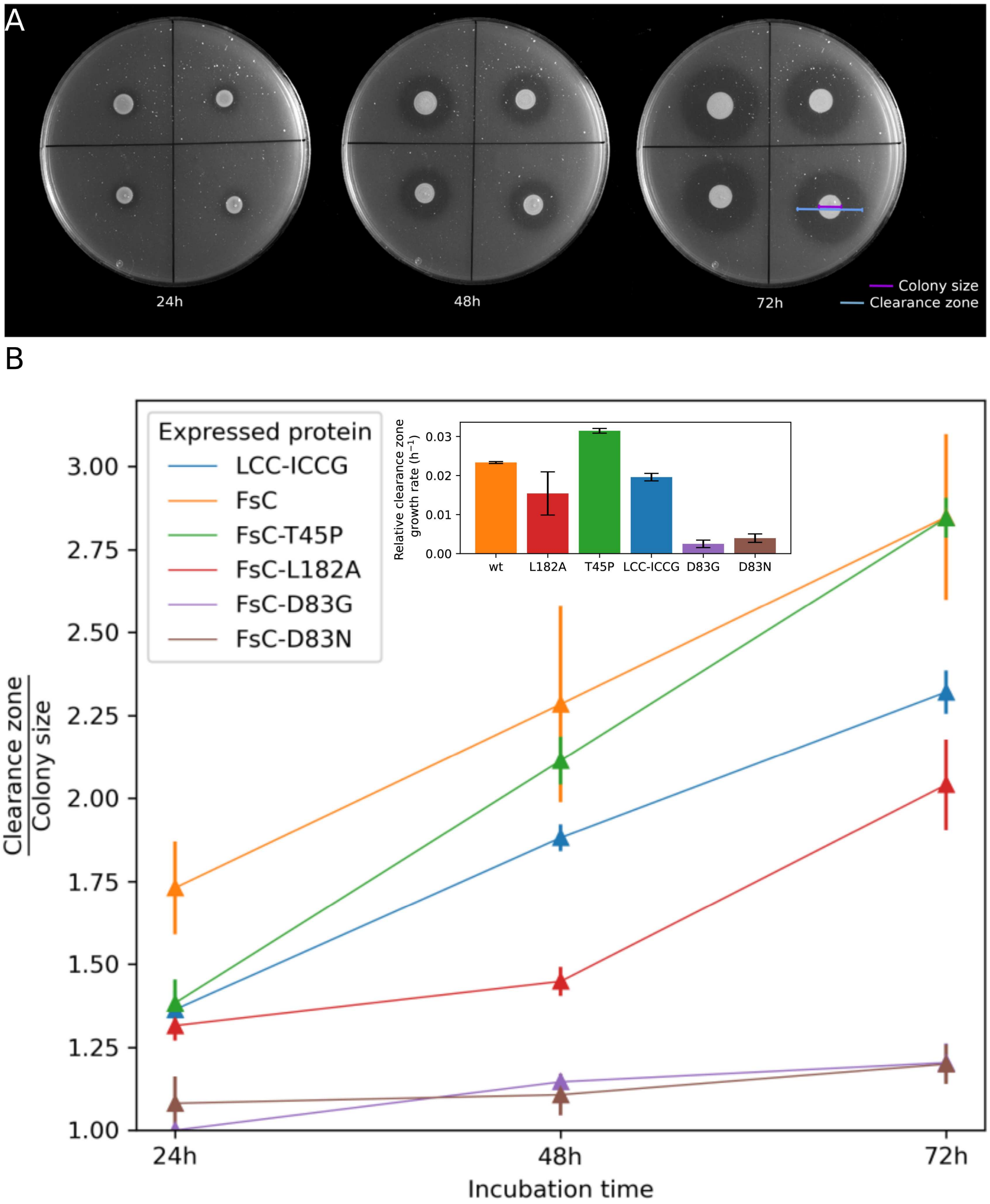


Figure S3: Quantitative plate-clearing assay of *V. natriegens* expressing LCC-ICCG and FsC enzymes (wt, T45P, L182A, D83G, D83N). All enzymes were tested in four replicates. A: The four replicates of FsC-T45P imaged at 24 h, 48 h, and 72 h. Colony diameter is indicated in purple; clearance diameter in blue. For reference, the plates are 90 mm wide. B: Ratios of clearance zones to colony sizes at 24 h, 48 h, and 72 h. Error bars represent the standard deviation of the mean for the four replicates of each enzyme. The insert shows the growth rate as the slope of a linear regression calculated from the average data points.

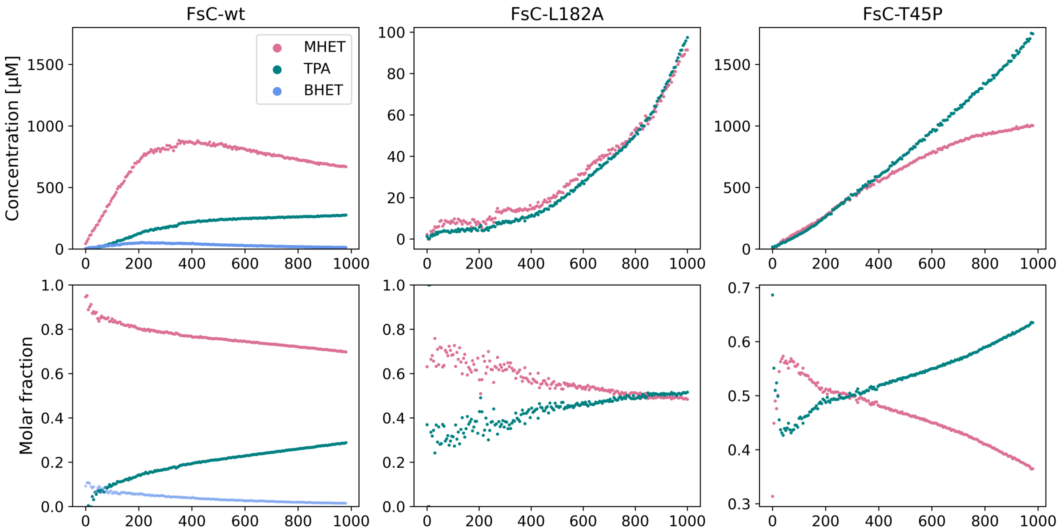


Figure S4: Time courses of PET hydrolysis by FsC-T45P measured by time-resolved $^1$H-NMR at 40°C. Data for FsC-wt and L182A are shown for comparison, reproduced with permission from Hellesnes, Vijayaraj et al. [1]. Spectra were recorded every 5 minutes for 16.3 hours. The top panels show BHET (blue), MHET (pink), and TPA (green) concentrations over time, calculated from their ratios relative to a standard TSP signal at 400 µM. The bottom panels show the molar fractions of BHET, MHET, and TPA over time.

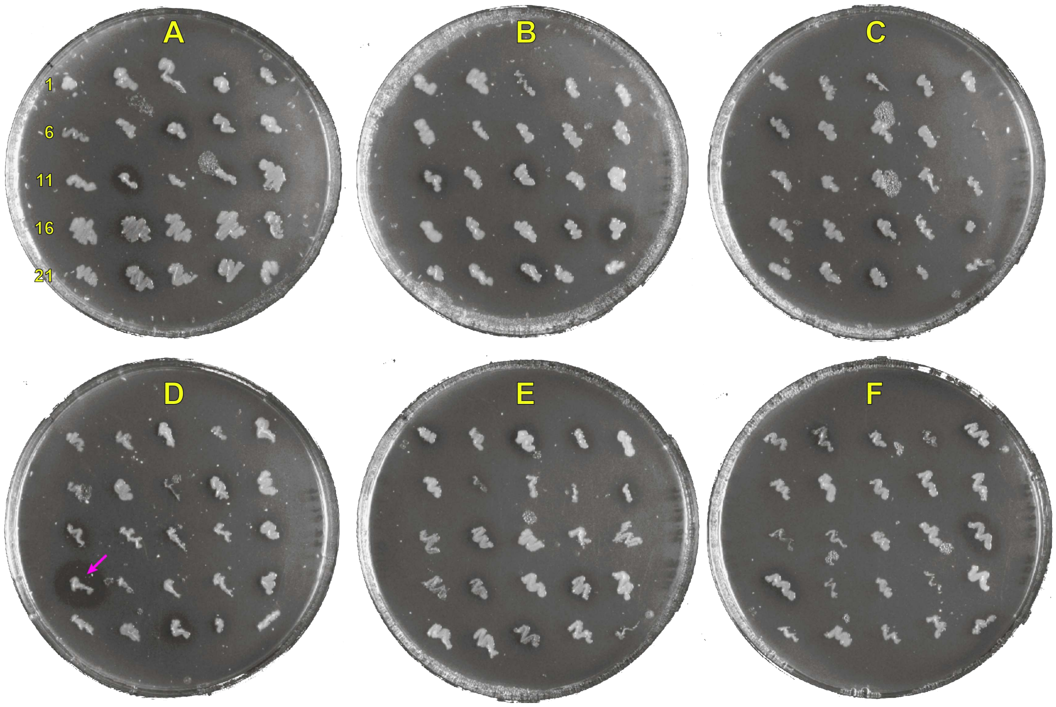


Figure S5: A photograph of six plates, each containing 25 clones of *V. natriegens* harboring plasmids from the FsC DNA library. Clearance zones around colonies indicate BHET hydrolytic activity. Colonies were assigned an identification from left to right and top to bottom (plates A to F; colonies 1 to 25) as indicated. This nomenclature is used in Table S1. Colony D16 (FsC-I24T) is marked with a pink arrow.

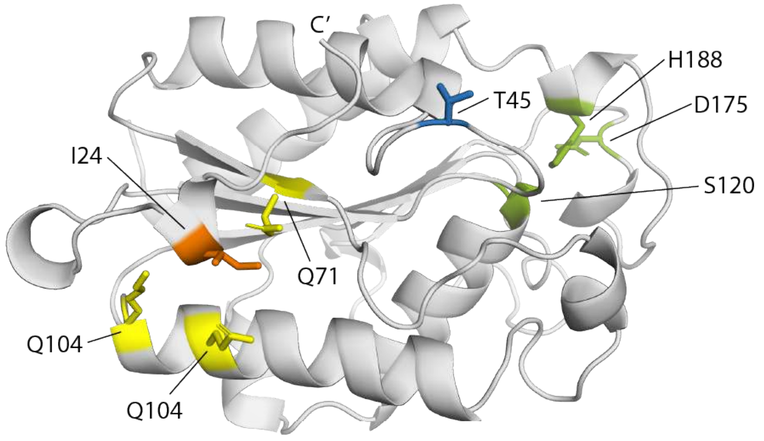


Figure S6: Crystal structure of FsC (PDB: 1CEX [2]), with the catalytic triad (green), T45 (blue), I24 (orange), and surrounding glutamine residues (yellow) shown as sticks.

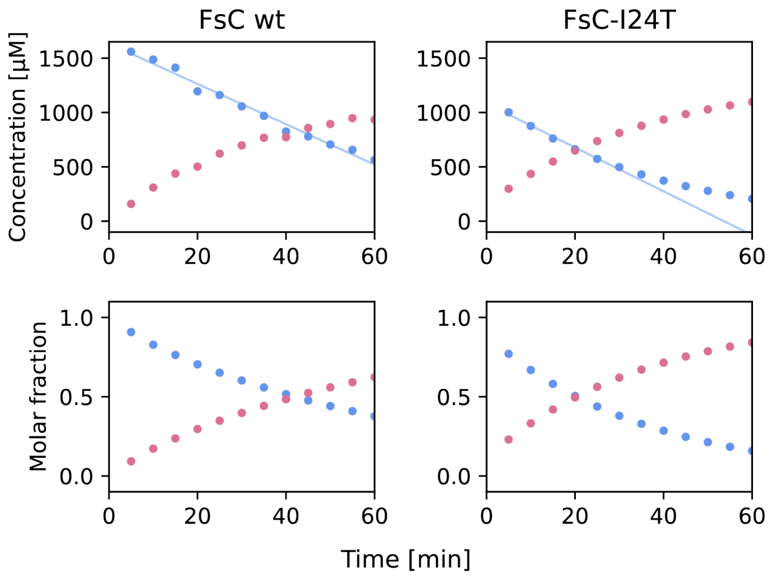


Figure S7: Time courses of BHET hydrolysis by FsC-wt and the FsC-I24T variant generated from error-prone PCR of the wild-type plasmid. The top panels depict the concentration (µM) of BHET (blue) and MHET (pink) over time, calculated from the ratio to a standard TSP signal at 400 µM. The bottom panels show the molar fractions of BHET and MHET over time. The initial BHET degradation rates (blue lines) were calculated to be 13.9 ± 1.2 (FsC-wt) and 15.1 ± 1.6 (FsC-I24T) µmol $g^{-1}$ enzyme $s^{-1}$.

Table S1: Substitutions identified in FsC variants from error-prone PCR, categorized as active or inactive. Substitutions are color-coded according to whether they are conservative (blue) or non-conservative (red) according to the BLOSUM 62 substitution matrix [3].

| **Variant** | **AA substitutions** | | | |
|---|---|---|---|---|
| Active variants | | | | |
| A4 | S30P | | | |
| A5 | N106D | | | |
| A9 | I159V | | | |
| A12 | D111G | R208L | | |
| A17 | V34A | | | |
| A22 | A29V | | | |
| B20 | S181N | | | |
| C3 | D134G | | | |
| D16 | I24T | | | |
| F5 | L114V | I137L | | |
| F16 | R166G | | | |
| Inactive variants | | | | |
| A6 | N5S | Y149C | K151R | G157C |
| B9 | S61P | C171S | D175G | |
| D1 | H188P | | | |
| E11 | E97G | Y162C | | |
